# Chromatin-associated effectors of energy-sensing pathways mediate intergenerational effects

**DOI:** 10.1101/2020.08.31.275727

**Authors:** Pedro Robles, Anisa Turner, Giusy Zuco, Panagiota Paganopolou, Beth Hill, Vikas Kache, Christine Bateson, Andre Pires-daSilva

## Abstract

Environmental stimuli experienced by the parental generation influence the phenotype of subsequent generations. The effects of these stimuli on the parental generation may be passed through the germline, but the mechanisms of this non-Mendelian type of inheritance are poorly known. Here we show that modulation of nutrient-sensing pathways in the parental generation of a nematode (*Auanema freiburgensis*) regulates phenotypic plasticity of its offspring. In response to pheromones, AMP-activated protein kinase (AMPK), mechanistic target of rapamycin complex 1 (mTORC1) and insulin signaling regulate stress resistance and sex determination across a generation. The effectors of these pathways are closely associated with the chromatin and their regulation affects the acetylation chromatin status in the germline. These results suggest that highly conserved metabolic sensors regulate phenotypic plasticity by changing the epigenetic status of the germline.

## INTRODUCTION

The phenotype of an individual is the result of the interactions between its genome and the environment. However, the phenotype may also be influenced by experiences of the parents: parental environment, such as diet, may result in epigenetic changes in the germline that cause non-adaptive phenotypes in the offspring (Chen et al., 2016, Sharma et al., 2016). An example case in humans suggests that famine increases the risk of metabolic defects in one or more generations (Kaati et al., 2007).

However, there are also mechanisms for passing information about the maternal environment to the offspring that increase fitness (Burton et al., 2017, Dantzer et al., 2013, Jablonka, 2013). This is referred to as adaptive phenotypic plasticity, which allows parents to match the phenotype of their offspring to changes in the local environment (West-Eberhard, 2003). For example, by sensing environmental cues, some animals can generate predator-resistant offspring (Agrawal et al., 1999, Gilbert, 2017), or stress-resistant offspring adapted to seasonal conditions (Mousseau and Dingle, 1991). Relatively little is known about mechanisms in which the parental generation senses the environment to induce adaptive phenotypic plasticity across one (intergenerational) or more generations (transgenerational) (Perez and Lehner, 2019).

Invertebrate model systems, such as the nematode *Caenorhabditis elegans*, have been instrumental in revealing some of the mechanisms of inter- and transgenerational inheritance (Miska and Ferguson-Smith, 2016, Perez and Lehner, 2019). The free-living nematode *Auanema freiburgensis* is an attractive new animal model system for studying the mechanisms of inheritance of parental effects (Kanzaki et al., 2017, Zuco et al., 2018, Anderson et al., 2020). This is because the assays for studying the mechanisms of inheritance of parental effects in *A. freiburgensis* are fast and easy to perform due its short generation time (~4 days at 20 °C) and easy-to-distinguish morphologies in the offspring.

*A. freiburgensis* produces three sexes, consisting of males, females and hermaphrodites (Kanzaki et al., 2017). The male of *A. freiburgensis* is determined genetically (XO), by mechanisms that will be addressed in a separate report. The hermaphrodite versus female sex (both XX) is determined by the environment experienced by the mother. Hermaphrodite individuals kept in isolation produce mostly female offspring, whereas hermaphrodites exposed to high population density conditions produce mostly hermaphrodite offspring (Fig. 1A).

**Figure 1.**
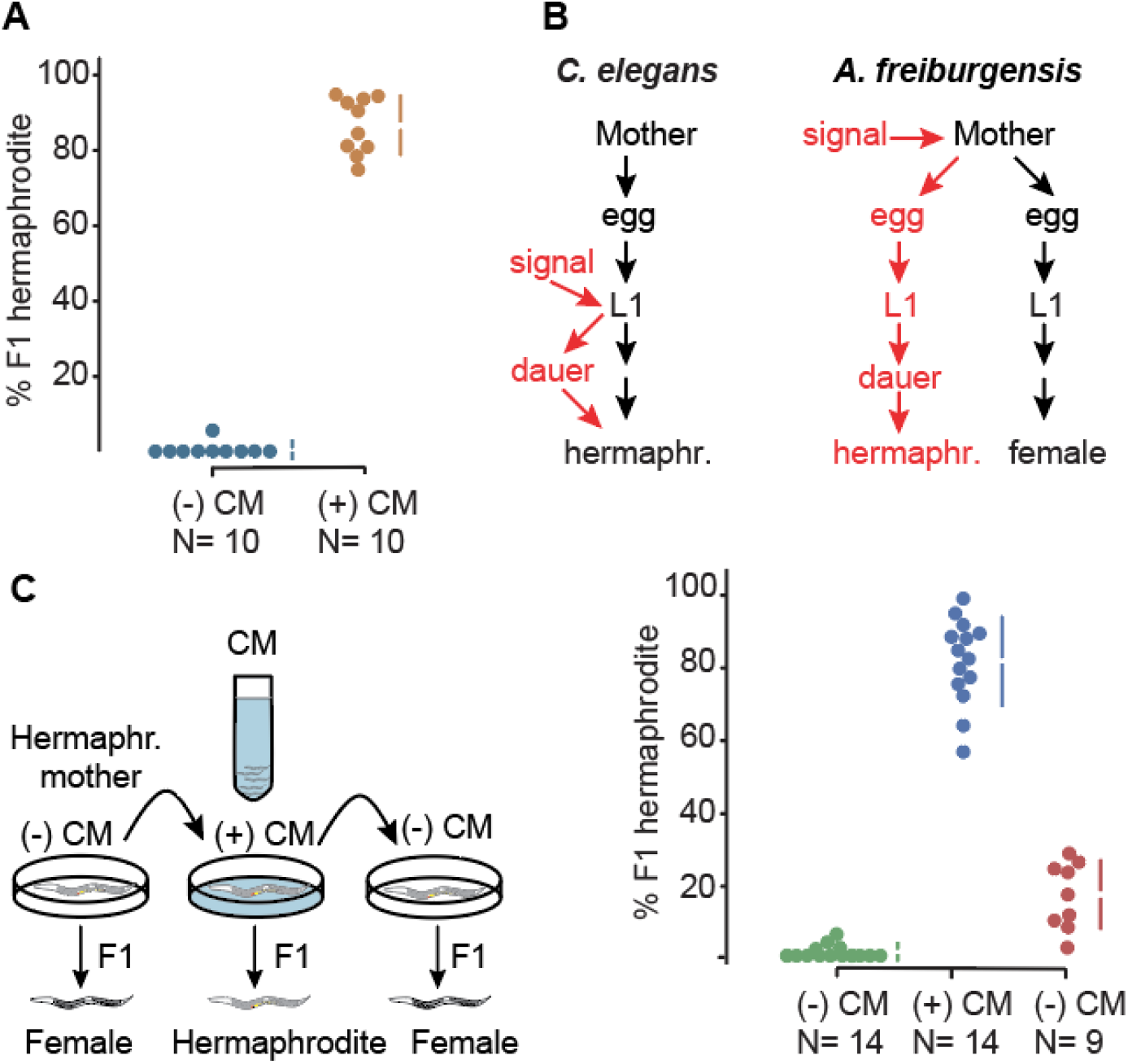
Dauer and hermaphrodite development are induced across generations in *A. freiburgensis*. **A.** When hermaphrodite mothers are cultured in non-crowding conditions ((-) CM), most of the XX F1s are female. (10 broods, from which a total of 149 F1s were scored). When mothers are in crowding conditions ((+) CM), most of the XX F1s are hermaphrodites (10 broods, with a total of 199 F1s scored). The data in colored dots represent the percentage of F1 hermaphrodites in each brood and is plotted on the upper axes. The colored vertical lines indicate ± SD and the mean is represented as a gap in the lines. N= sample size. **B.** In *C. elegans*, the L1 larvae respond to environmental signals to facultatively form stress-resistant dauers. In *A. freiburgensis*, it is the mother and not the L1s that respond to environmental signals. *A. freiburgensis* dauers obligatorily develop into hermaphrodite adults. **C.** In the experimental setup (top), the same individual mother hermaphrodite was transferred every 24 hours to a new environmental condition. Initially, it was placed in a plate without conditioned medium (-) CM, followed by the transfer to a (+) CM plate and then to a new (-) CM plate. The plot representation is the same as for Fig. 1A. On the last day, 5 mothers died and thus only 9 broods were scored.

Here we show that high-density population conditions experienced by the *A. freiburgensis* mother, a signal for imminent starvation, triggers the formation of F1 dauer larvae. These dauers develop into hermaphrodite adults, while non-dauer larvae develop into female adults (or males). Pharmacological assays indicate that energy-sensing signaling mediated by AMP-activated protein kinase (AMPK), mechanistic target of rapamycin complex 1 (mTORC1), and insulin signaling are involved in intergenerational inheritance in *A. freiburgensis*. Effectors of these pathways are associated with chromatin, which changes the histone acetylation status in the germline chromatin to produce F1 dauers, which then develop into hermaphrodite adults.

## RESULTS

A crucial factor in the development of *Auanema* hermaphrodites is the passage through the stress-resistant dauer stage (Félix, 2004, Chaudhuri et al., 2011, Kanzaki et al., 2017, Chaudhuri et al., 2015), which has morphological and behavioral adaptations for dispersal. In *A. freiburgensis*, all XX larvae that pass through the dauer stage become hermaphrodites (N= 96), whereas XX non-dauer larvae develop into females (N= 93). Similar to *A. rhodensis* (Chaudhuri et al., 2011) and other trioecious nematodes (Johnigk and Ehlers, 1999), we never observed *A. freiburgensis* males to undergo dauer formation. Thus, environmental stressors experienced by the maternal generation of *A. freiburgensis* are used as a signal to generate non-feeding offspring that can survive starvation conditions and reproduce by self-fertilization once food becomes available. In summary, these results suggest that dauer formation in *A. freiburgensis* is induced across a generation, instead of within the same generation as in *Caenorhabditis elegans* (Cassada and Russell, 1975) (Figure 1B).

High population density conditions were induced by incubating *A. freiburgensis* hermaphrodites with conditioned medium (CM) derived from liquid cultures containing high nematode population densities (see Methods). Importantly, only the parental generation was exposed to the CM. The induction of dauers through the hermaphrodite mother is limited to one generation: F1 hermaphrodites derived from mothers in (+) CM plates produce mostly female offspring (99.6% out of 470 F2 offspring, scored from 10 broods). To test if *A. freiburgensis* L1 larvae can also respond to crowding conditions, eggs derived from mothers cultured in isolation were left to hatch and undergo larval development in (+) CM plates until adulthood. 95.7% (N= 161) of these L1s developed into females, indicating that larvae do not respond to crowding conditions. To investigate if other maternal environmental conditions affect the sexual fate of the F1s, mothers were incubated for 24-hours to high temperature (25 °C) or starvation. Most XX offspring (97%) developed into female adults for both conditions (166 F1s scored from mothers at 25 °C and 146 F1s scored from starving mothers). These results indicate that the conditioned medium is the only environmental stressor that induces intergenerational polyphenism in *A. freiburgensis* on its own.

To test the minimal population density sufficient for the induction of dauers and hermaphrodites across a generation, we incubated the maternal generation at different densities. When cultured for 6 hours, a minimum density of 16 adult hermaphrodites per cm^2^ is sufficient for the induction of 100% (N= 295 F1s) of hermaphrodite offspring. In densities below 10 individuals/cm^2^, the hermaphrodite mothers produce only female offspring (10 individuals/cm^2^: 100% females, N= 78 F1s; 6 individuals/cm^2^: 98.5% F1 female, N= 66 F1s). At an intermediate density (13 individuals/cm^2^), hermaphrodites produce 19% (N= 126 F1s) of hermaphrodite offspring.

### Modulation of AMPK signaling changes hermaphrodite/female sex ratios in *A. freiburgensis*

In eukaryotes, caloric restriction triggers the activation of AMP-activated protein kinase (AMPK) (Apfeld et al., 2004), a highly conserved energy sensor (Hardie et al., 2012). AMPK activity protects cells against the depletion of ATP by stimulating energy-producing pathways and inhibiting energy-consuming processes (Carling, 2004). In *C. elegans*, AMPK is required for lifespan extension and germline viability when the nematode is in nutrient stress (Apfeld et al., 2004, Narbonne and Roy, 2006, Fukuyama et al., 2012, Demoinet et al., 2017). The full kinase activity of AMPK requires phosphorylation of threonine residue 172 (Thr172) by upstream kinases (Stein et al., 2000, Lee et al., 2008, Apfeld et al., 2004).

Since high population density is likely to result in imminent food scarcity, we reasoned that the AMPK pathway may be involved in intergenerational inheritance in *A. freiburgensis*. High population density conditions were induced by incubating *A. freiburgensis* hermaphrodites with conditioned medium (CM) of high population density liquid cultures (see Methods). We hypothesized that AMPK regulates target proteins in the maternal germline to influence the phenotype of the following generation. To test this hypothesis, we first tested the levels of an enzyme that activates AMPK, Liver Kinase B1 (LKB1). LKB1, known in *C. elegans* as PAR-4 (Watts et al., 2000, Lee et al., 2008), phosphorylates and activates AMPK in the context of energy stress (Woods et al., 2003, Hawley et al., 2003). LKB1 is part of a complex with two proteins Ste20-related adaptor protein-alpha (STRAD) (Baas et al., 2003) and mouse protein 25-alpha (MO25alpha) (Boudeau et al., 2003). Antibody staining against LKB1 and STRAD showed a higher level of staining in the meiotic portion of the germline isolated from animals cultured in crowding conditions (Fig. 2A, C, Supplemental Figure 1). Their localization was predominant in the cytoplasm of germline cells (Supplemental Figure 1).

**Figure 2.**
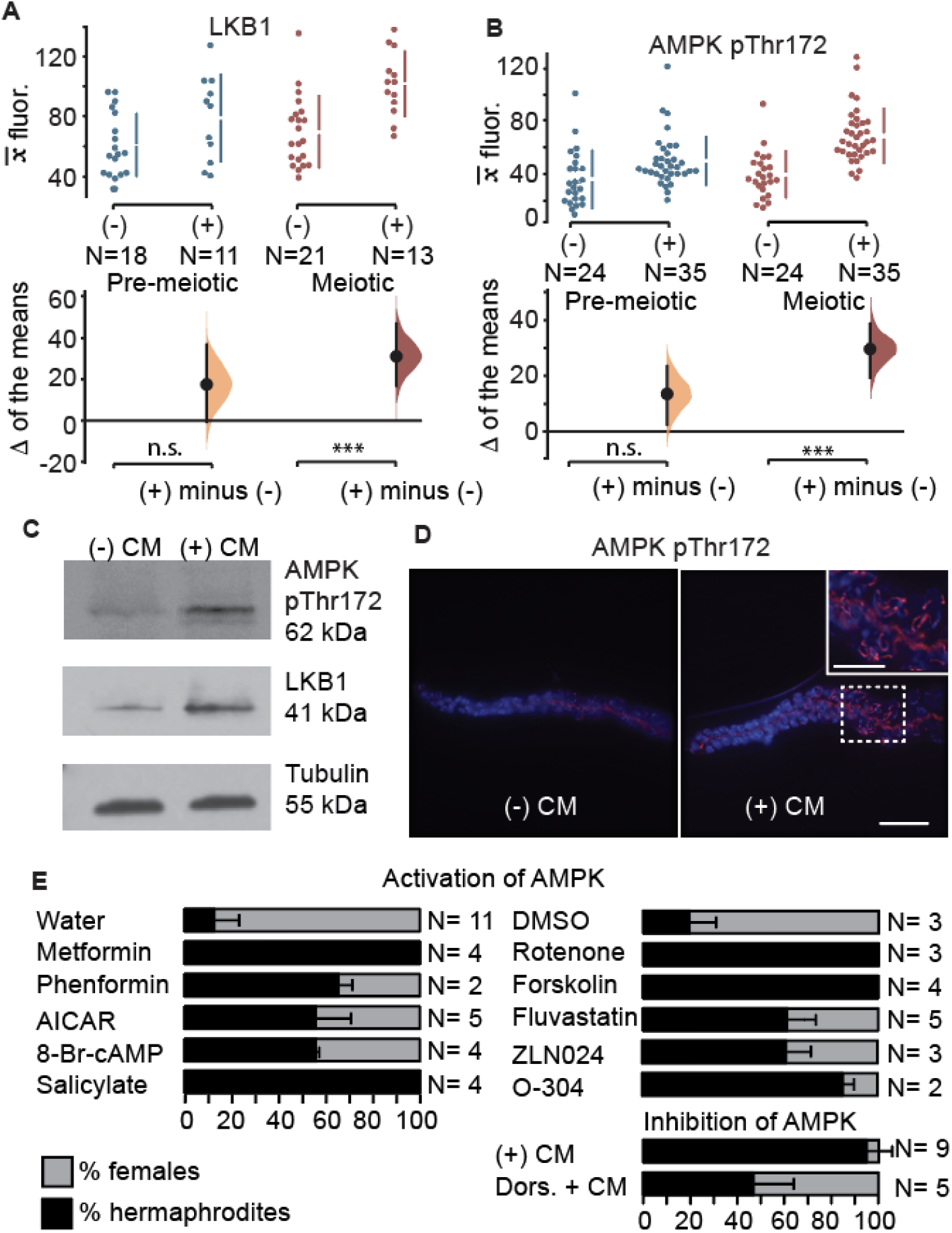
AMPK pathway modulation in the *A. freiburgensis* germline. (**A, B**) Mean antibody fluorescence (*x__*) in the pre-meiotic (blue) and meiotic portion (red) of the germline, in the absence (-) or presence (+) of conditioned medium. N= sample sizes. The mean difference for the two comparisons is shown as a Gardman-Altman estimation plot. The raw data is plotted on the upper axes, with colored vertical lines indicating ± 95% CI, and the mean is represented as a gap in the lines. Each difference of the means is plotted on the lower axes as a bootstrap sampling distribution. The difference of the means is depicted as a black dot and 95% confidence intervals are indicated by the black vertical error bars. n.s., p>0.05; ***= p ≤ 0.001. (**C**) Western blots with proteins derived from hermaphrodites incubated in the absence (-) CM or presence (+) CM of conditioned medium. (**D**) AMPK pThr172 antibody staining of gonads dissected from hermaphrodites incubated in either (-) CM or (+) CM. Bar, 15 μm. Insert in the right picture is a magnification from the region marked with a stippled square. Bar, 7.5 μm. (**E**) Mean percentage and SD of hermaphrodite and female F1 offspring from hermaphrodites treated with chemicals. The control was either using water (left) or DMSO (right), depending on how the chemical compounds were dissolved. Dors.= Dorsomorphin. In all cases, diluted (1:10) CM was added to the medium, with exception to plates with dorsomorphin, which had undiluted CM. N= number of replicates.

To test the levels of AMPK, we used an antibody that detects the active, phosphorylated form of AMPK (AMPK pThr172). Consistent with the higher levels of LKB1 and STRAD, we also found that the anti-AMPK pThr172 staining was stronger in crowding conditions compared to control animals (Fig. 2 B-D). The difference in the level of staining was restricted to the meiotic region of the germline (Fig. 2, D) and the AMPK staining is closely associated with the chromatin of pachytene cells (Fig. 2D).

To functionally test the role of AMPK in mediating intergenerational inheritance in *A. freiburgensis*, we used pharmacological compounds that modulate AMPK activity. We measured the effects of these compounds on intergenerational inheritance by scoring hermaphrodite and female sexes in the offspring. As mentioned previously, high population densities induce the production of dauer larvae in the F1, which mature to become hermaphrodite adults. Consistent with a role of AMPK in mediating this effect, we found that AMPK activators induce the production of hermaphrodites (Fig. 2E) (for a recent review on pharmacological activation of AMPK, see (Steinberg and Carling, 2019)). In most cases, these compounds cause changes in the F1 sex ratios when on their own (Supplemental Figure 2), but potentiation of their effects was significantly stronger when diluted CM (1:10 CM) was added to the culture medium. This may indicate that synergistic effects of different mechanisms are necessary to fully elicit a robust response, or that those energy-sensing pathways can be efficiently activated only when upstream events occur first.

Although the mechanisms of action are not clear for all pharmacological compounds, they can be broadly divided into indirect and direct AMPK activators. Any treatments that raise the AMP/ADP:ATP ratios are expected to indirectly activate AMPK. For instance, inhibition of mitochondrial respiration by metformin, phenformin and rotenone have been implicated in the activation of AMPK (El-Mir et al., 2000, Owen et al., 2000, Zhou et al., 2001, Sakamoto et al., 2004, Shaw et al., 2004, Huang et al., 2008, Toyama et al., 2016, Hou et al., 2018). Forskolin, an adenylate cyclase activator, activates AMPK by increasing the cytosolic cAMP concentration (Seamon et al., 1983). Statins, such as fluvastatin (Xenos et al., 2005), have been proposed to activate AMPK. The incubation of mothers with all these compounds resulted in a higher proportion of hermaphrodite progeny (Fig. 2E).

Compounds that are similar to AMP can activate AMPK directly. 5-Aminoimidazole-4-carboxamide ribonucleotide (AICAR), for example, increases the activity of AMPK after being converted to an AMP analog inside the cell (Corton et al., 1995), whereas 8-Br-cAMP is a non-hydrolyzable analog of cAMP (Hussey et al., 2017). Other compounds, such as the plant product salicylate (Hawley et al., 2012), and the synthetic compounds ZLN024 (Zhang et al., 2013) and O-304 (Steneberg et al., 2018) bind to AMPK, causing allosteric activation and inhibition of dephosphorylation of the pThr172. All these compounds induced a higher percentage of hermaphrodite offspring than controls (Fig. 2E). To inhibit AMPK, we used dorsomorphin (Zhou et al., 2001). As expected, hermaphrodites in CM with dorsomorphin resulted in a lower proportion of hermaphrodite progeny compared to controls (Fig. 2E).

### Maternal inhibition of mTORC1 signaling results in mostly hermaphrodite offspring

Since energy-sensing by AMPK induced intergenerational effects in *A. freiburgensis*, we hypothesized that other energy sensors may be involved in the same process. The intracellular nutrient sensor mTORC1 complex is a multisubunit kinase that senses growth signals and stimulates anabolism when nutrients are abundant (Kapahi et al., 2010, Ma and Blenis, 2009, Wullschleger et al., 2006, Zoncu et al., 2011, Laplante and Sabatini, 2012). Therefore, we would predict that in low population densities and readily available nutrients, the mTOR pathway would be active in *A. freiburgensis*. Under these conditions, *A. freiburgensis* produces mostly non-dauer larvae that later become female offspring. To investigate the kinase activity of mTORC1, we examined the expression of a well-characterized target protein, p70 S6K protein kinase (S6K) (Kapahi et al., 2010). Antibody staining against the phosphorylated form of S6K (S6K pThr389) was detected primarily in germline cells isolated from animals grown in low-density conditions (Fig. 3A-C). Most staining was associated with the chromatin, both in mitotic cells (Fig. 3A) and meiotic cells in late pachytene stages (Fig. 3B). Since AMPK and mTORC1 have opposing actions (Hindupur et al., 2015), we hypothesized that treatment of animals with metformin, an activator of AMPK, would inhibit mTORC1 signaling. Consistent with this hypothesis, we found that treatment of animals with metformin resulted in a smaller number of cells stained with SK6 pThr389 (Fig. 3D).

**Figure 3.**
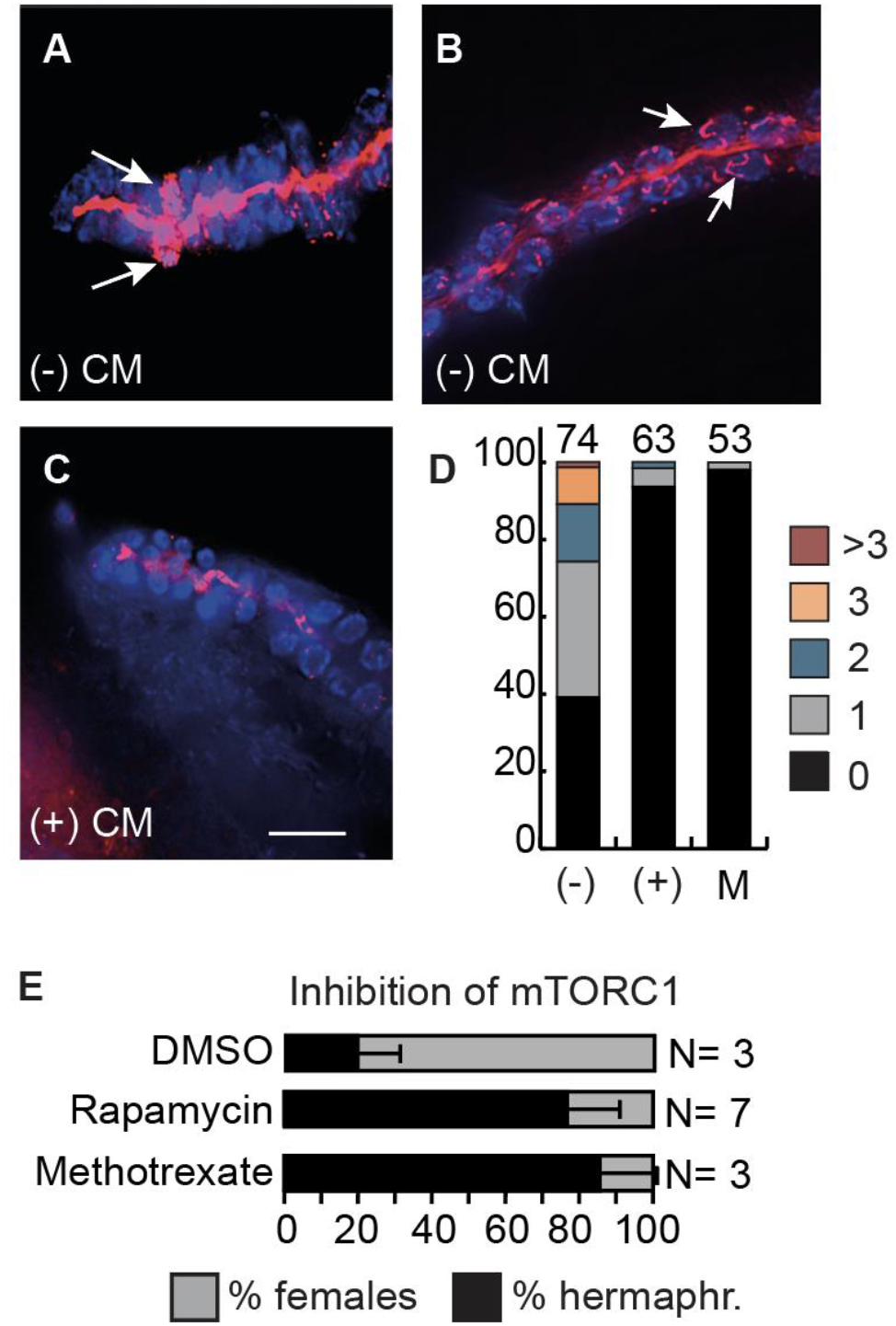
mTOR signaling modulates intergenerational inheritance in *A. freiburgensis*. (**A-C**) Staining with S6K pThr389 antibody (red) and DAPI (blue) of dissected gonads from hermaphrodites incubated either in the absence ((-) CM) or in the presence ((+) CM) of conditioned medium. The arrows indicate cells marked with the antibody in the premeiotic region (**A**) and in the meiotic region (**B**), respectively. In (**C**), only the rachis has staining. Bar, 15 μm. (**D**) Percentages of gonads with signal for S6K pThr389 antibody staining. The different colors represent the percentage of gonads with at 0, 1, 2, 3 or more than 3 cells stained in the premeiotic (PM) tip. Quantification was performed from gonads isolated from animals in the absence (-) or in the presence (+) of conditioned medium, and in the presence of metformin (M). The number of gonads analyzed is indicated on the top of the bars. (**E**) Mean percentage and SD of hermaphrodite and female F1 offspring from hermaphrodites incubated with either DMSO or pharmacological compounds, together with some CM (1:10) CM. N= number of replicates.

To test the effect of modulating mTORC1 activity on sex ratios, we treated mothers with pharmacological compounds. Mothers treated with rapamycin (Heitman et al., 1991, Robida-Stubbs et al., 2012) produced a greater proportion of F1 hermaphrodites than control mothers (Fig. 3E). mTORC1 signaling promotes nucleic acid synthesis, as long as nucleotide precursors are available (Hoxhaj et al., 2017). Treatment with methotrexate, a chemical that suppresses the *de novo* purine synthesis enzymes (Rajagopalan et al., 2002), inhibits mTORC1 activity. We found that *A. freiburgensis* hermaphrodites treated with methotrexate generated mostly hermaphrodite offspring (Fig. 3E). Altogether, these results indicate that mTOR signaling is involved in intergenerational inheritance in *A. freiburgensis*.

### Insulin signaling is downregulated in animals in crowding conditions

The insulin signaling pathway regulates metabolism, development, and lifespan in a wide variety of animals. One of the regulators of the insulin pathway is a conserved phosphatase named PTEN (or DAF-18 in *C. elegans*)(Solari et al., 2005). To examine the regulation of the insulin pathway in *A. freiburgensis*, we used an antibody against PTEN/DAF-18 to stain isolated gonads from hermaphrodites cultured in low- and high-density conditions. We found that the antibody against PTEN/DAF-18 stained more strongly the germline when hermaphrodites were incubated in high-density conditions than in low-density populations (Fig. 4A, Supplemental Figure 3).

**Figure 4.**
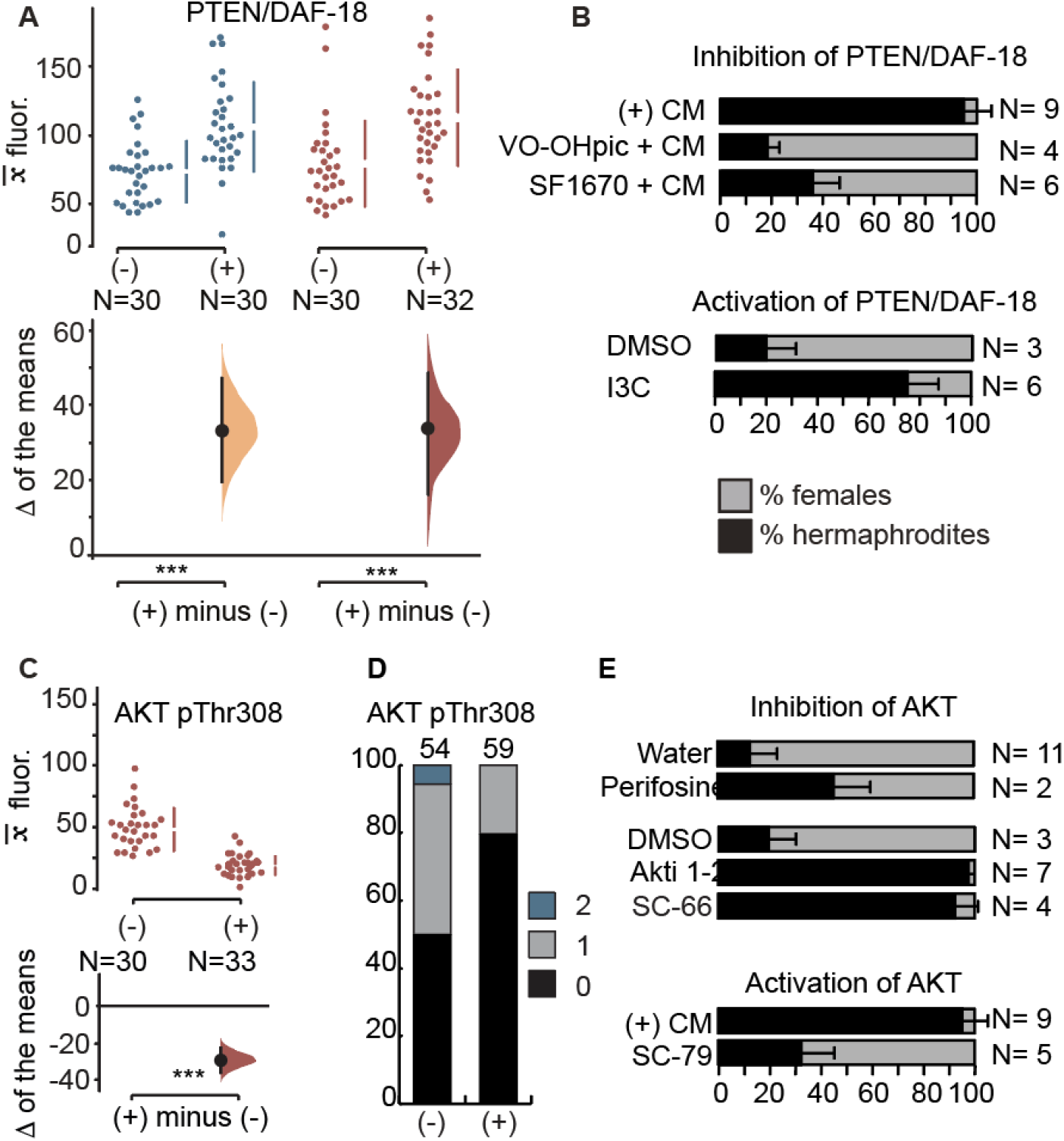
Regulation of PTEN/DAF-18 and AKT. (**A**) Quantification of antibody staining with PTEN/DAF-18 in the maternal gonads. The representation and labeling of graphs are as in Fig. 2A. (**B**) Mean percentage and SD of hermaphrodite and female F1 offspring from hermaphrodites treated with chemicals that activate or inhibit PTEN/DAF-18. (+) CM represents undiluted conditioned medium. The DMSO control and Indole-3-Carbinol (I3C) incubations were performed with diluted (1:10) CM. N= number of replicates. (**C**) Quantification of antibody staining for AKT pThr308 in the meiotic portion of the germline, with representation as in (**B**). (**D**) Quantification of meiotic germline cells with staining with an antibody against AKT pThr308, with graphical representation as in Fig. 3D. (**E**) Effect of pharmacological inhibition or activation of AKT on sex ratios in the F1s.

To test if PTEN/DAF-18 mediates the generation of hermaphrodites, we used the PTEN/DAF-18 inhibitors VO-OHpic (Rosivatz et al., 2006) and SF1670 (Li et al., 2011). When in the presence of conditioned medium from high-density populations, hermaphrodites treated with those inhibitors generated mostly female offspring (Fig. 4B). Activation of PTEN/DAF-18 in hermaphrodites with the compound Indole-3-Carbinol under low population densities resulted in mostly hermaphrodites (Fig. 4B).

One of the target proteins and effectors for insulin signaling is AKT kinase (also known as PKB) (Paradis and Ruvkun, 1998), which among several anabolic functions, also prevents chromatin condensation (Martelli et al., 2012, Manning and Cantley, 2007, Manning and Toker, 2017). Maximal activation of AKT requires phosphorylation at residues Thr308 and Ser473 (Alessi et al., 1996). Immunostaining with antibodies against AKT pThr308 (Fig. 4C, D, Supplemental Figure 4) revealed that staining is prominently associated with the chromatin in germline cells of animals grown under non-crowding conditions. No such association is seen when animals are in crowding conditions (Fig. 4C). The same pattern is seen for AKT pSer473 (Supplemental Figure 4). Maternal inhibition of AKT with the chemicals perifosine (prevents activation of AKT by affecting its subcellular localization) (Kondapaka et al., 2003), Akti-1/2 (stabilizes the inactive conformation of AKT) (Barnett et al., 2005), and SC-66 (allosteric inhibitor of AKT) (Jo et al., 2011) results in a higher proportion of hermaphrodite progeny (Fig. 4E). On the other hand, activation of AKT with SC-79 (Jo et al., 2012) prevents the generation of hermaphrodite progeny when the mother is in crowding conditions (Fig. 4E). Altogether, these results are consistent with the hypothesis that crowding conditions induce a lower insulin signaling, causing the production of hermaphrodite offspring.

### Changes in the maternal histone acetylation status modulate sex ratios in the F1

Energy-sensing pathways have been implicated in the regulation of acetylation of histones, histone modifiers, and cellular proteins (Salminen et al., 2016). To examine if acetylation patterns change in the germline when *A. freiburgensis* is in high population densities, we compared the level of antibody staining in gonads isolated from hermaphrodites cultured in the absence or presence of CM. Antibody staining against acetylated residues on histones 3 and 4 was at higher levels in the germline derived from animals cultured in the presence of CM compared to controls, both for premeiotic and meiotic portions (Fig. 5A-B, Supplemental Figure 5). The same trend was observed when using an antibody that binds to all acetylated proteins (pan-LysAc), although differences were detected only for the premeiotic portion of the germline (Fig. 5C).

**Figure 5.**
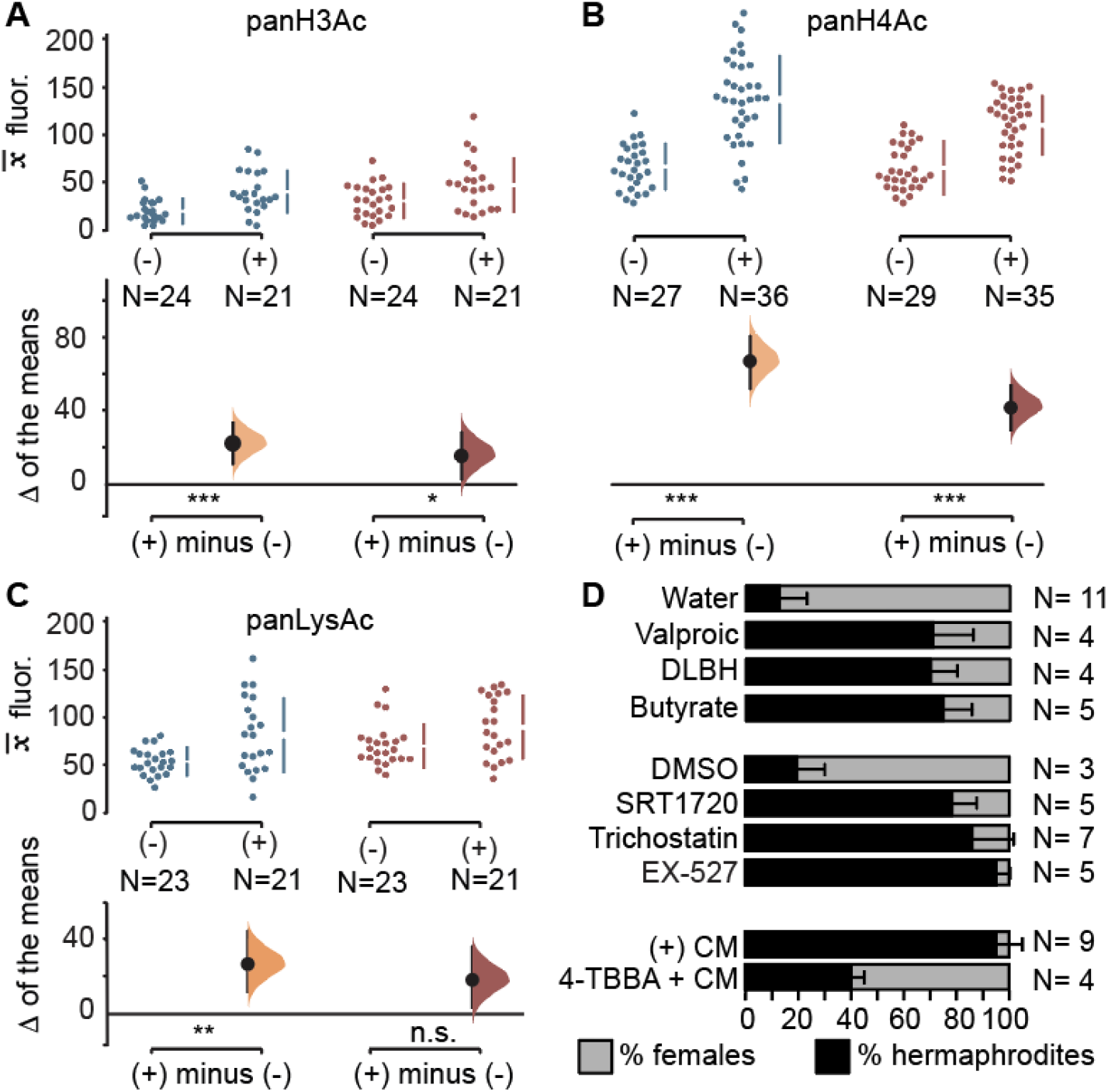
Regulation of acetylation levels. Mean antibody fluorescence (*x__*) for panH3Ac (**A**), panH4Ac (**B**) and panLysAc (**C**) in the pre-meiotic (blue) and meiotic portion (red) of the germline, in the absence (-) or presence (+) of conditioned medium. N= sample sizes. The P values are calculated from a Mann-Whitney test (U): n.s., p>0.05; * = p ≤ 0.05; ** = p ≤ 0.01; ***= p ≤ 0.001. (**D**) Mean percentage and SD of hermaphrodite and female F1 offspring from hermaphrodites treated with chemicals. DLBH: DL-β hydroxybutyrate; TBBA: 4-tert-butylbenzoic acid. (1:10) CM was added to the medium, except plates with 4-TBBA, which had undiluted CM. N= number of replicates.

To test if modulation of acetylation levels causes changes in sex ratios, we induced hyperacetylation by treating *A. freiburgensis* hermaphrodites with the histone deacetylase inhibitors SRT1720 (Zarse et al., 2010, Milne et al., 2007), Trichostatin A (Yoshida et al., 1990), Valproic Acid (Evason et al., 2008, Forthun et al., 2012), D-β-hydroxybutyrate (Edwards et al., 2014), Butyrate (Zhang et al., 2009) and EX-527 (Solomon et al., 2006). In all cases, more hermaphrodites than females were produced relative to control (Fig. 5D). By contrast, incubating the mothers in high-density conditions together with the inhibitor of acetylation 4-tert-butylbenzoic acid (Chen et al., 2014) resulted in less hermaphrodite offspring (Fig. 5D).

## DISCUSSION

*Auanema* nematodes have been isolated from similar environments as *C. elegans* (Félix and Duveau, 2012), which consists of ephemeral habitats with microbe-rich organic decomposing matter (Schulenburg and Felix, 2017, Kanzaki et al., 2017). Due to rapid population growth and quick depletion of resources, the ecology of these nematodes is characterized by a boom and bust population dynamics. In contrast to *C. elegans*, the developmental and phenotypic response to stress in *A. freiburgensis* occurs across a generation instead of within the same generation: maternal sensing of pheromones secreted by conspecifics induces the production of stress- and starvation-resistant dauer larvae. This indicates that the *A. freiburgensis* mother can predict the environmental conditions to which the offspring is likely to be exposed, and adjusts the F1 phenotype (dauer larvae) to temporarily survive in the absence of food. The *Auanema* dauers have migratory behaviors and always develop into selfing hermaphrodites (Kanzaki et al., 2017). By producing dauers that develop into hermaphrodites, a new population can be established even when the colonizing event is mediated by a single individual (Baker, 1955). This type of intergenerational inheritance, in which parental effects increase the fitness of both offspring and parents, has hallmarks for being adaptive (Uller, 2008).

Here we show that activators of AMPK and insulin signaling activators or mTORC1 inhibitors can mimic the exposure of *A. freiburgensis* to pheromones. These results indicate that highly conserved energy-sensing pathways are involved in mediating intergenerational inheritance in *A. freiburgensis* to generate stress-resistant offspring. How exactly do these energy-sensing pathways regulate phenotypic plasticity in the F1s? One possible mechanism is the direct regulation of the chromatin status in the maternal germline by the energy-sensing enzymes. Thus, activation of transcription of specific genes in the germline may determine the phenotype of the following generation. AMPK, for instance, has been shown to phosphorylate histones, which results in the activation of transcription (Bungard et al., 2010). Consistent with this, we found that protein levels increase for the activated form of AMPK when *A. freiburgensis* is under crowding conditions and is detected in close association with the chromatin of germline cells. By phosphorylating histones, AMPK has been shown to facilitate histone acetylation (Lo et al., 2001), thus promoting the transcription of a new set of genes (Lee et al., 1993).

Alternatively, AMPK may indirectly influence the chromatin status via activation of histone acetyltransferases (HATs) or inactivation of histone deacetyltransferases (HDAC), as demonstrated for other model systems (Shimazu et al., 2013, Yang et al., 2001). In *A. freiburgensis*, higher acetylation levels in the chromatin of the germline induced by crowding conditions results in stress-resistant offspring (Fig. 5). It remains to be established whether these acetylation levels are the result of direct phosphorylation of HATs and HDACs by AMPK, or indirectly by natural metabolites. As we show in Fig. 5D, natural metabolites indicative of metabolic stress that inhibit deacetyltransferases, such as D-β-hydroxybutyrate and butyrate (Shimazu et al., 2013), induce the production of stress-resistant offspring.

The strongest responses to the pharmacological compounds for the production of hermaphrodite progeny occurred when the animals were concomitantly exposed to diluted CM. In the complete absence of CM, only a few compounds elicited a strong response. This may indicate that pheromones in the CM activate more than one pathway and that they have to act in combination to elicit the full effect. Our findings that several energy-sensing pathways are involved in this process in *A. freiburgensis*, and that AMPK, insulin and TOR pathways are cross-regulated, are indicative of this hypothesis (González et al., 2020, Ruderman et al., 2010, Banerjee et al., 2016).

The concentration of the compounds used in our studies are relatively high compared to the ones used in mammalian cells (Burns et al., 2006). This is because nematodes have several physical and physiological adaptations that counteract xenobiotic agents (Burns et al., 2010). Like all pharmacological approaches, interpretation of the results must take into consideration possible lack of specificity (Corton et al., 1995, Longnus et al., 2003, Bain et al., 2007, Pacholec et al., 2010). To ameliorate the possibility of lack of specificity for AMPK activation, for instance, we used compounds that act through several mechanisms (high production of AMP, allosteric binding, protection against dephosphorylation, activation of phosphorylation). Genetic approaches using loss- and gain-of-function mutants will help to address some of the above-mentioned concerns (Adams et al., 2019).

As far as we know, the association of activated AMPK and S6K with the chromatin of germline cells has not been established in other organisms. The presence of AKT in the nucleus of germline cells may be associated with chromatin condensation, which would be reflected in transcription rates (Martelli et al., 2012). Our results indicate that these energy-sensing effectors acquired a new role in intergenerational inheritance in *A. freiburgensis* to regulate gene expression that influences the phenotype of subsequent generations. Given that AMPK, TOR, and insulin pathways are highly conserved in evolution, it is possible that they also mediate non-genetic inheritance via the germline in other organisms in which diet plays a role in determining phenotypic plasticity. Although the epidemiological studies in humans are indicative of diet playing such a role, the mechanisms for this are unknown (Horsthemke, 2018). The findings in this study provide the basis to test such a hypothesis.

## ACKNOWLEDGMENTS

This work was supported by Leverhulme Trust (RPG-2019-329). A.T. was funded by the Doctoral Training Program from Natural Environment Research Council (NERC CENTA) and P. P. was funded by the Doctoral Training from BBSRC (Midlands Integrative Biosciences Training Partnership). The authors would like to acknowledge the help of the Media Preparation Facility in The School of Life Sciences at the University of Warwick. Bacteria strains were provided by the CGC, which is funded by NIH Office of Research Infrastructure Programs (P40 OD010440).

## AUTHOR CONTRIBUTIONS

P.R., G.Z., V.K., and A.P.-d.S. designed the study. P.R., A.T., G.Z, V.K., P.P., B.H., C.B. and B.H. conducted experiments that involved the production of conditioned medium, incubation with chemicals, and sexing offspring. P.R. performed the immunocytochemistry experiments and P. P. performed the Western blots. P.R. and A.P.-d.S. wrote the paper.

## AUTHOR INFORMATION

The authors declare no competing interests.

## MATERIAL AND METHODS

### Strain and culture

We used the *Caenorhabditis elegans* N2 strain, and the *Auanema freiburgensis* strains SB372 (Kanzaki et al., 2017) and JU1782. The *A. freiburgensis* JU1782 strain was isolated from rotting *Petasites* stems sampled in Ivry, Val-de-Marne, France, in September 2009 by Marie-Anne Félix. Nematodes were cultured at 20 °C on standard Nematode Growth Medium (NGM) (Stiernagle, 2006) plates seeded with *Escherichia coli* OP50-1 strain. NGM medium was supplemented with 25 μg/mL nystatin and 50 μg/mL streptomycin to prevent microbial contamination.

### Sexing of progeny

To synchronize the age of the mothers, we collected dauers. *A. freiburgensis* dauers develop into hermaphrodite adults within 24 hours at 20 °C (Kanzaki et al., 2017). Dauer larvae are easily identified by their darker intestine and thinner body compared to similar-sized L3 larvae (which develop into females). Each dauer larva was placed on a 6 cm seeded NGM plate and incubated at 20 °C to develop into adulthood. Each egg laid by the parental (P0) generation was placed into single wells of a 96-well microtiter plate. After 3-5 days, the F1 was scored for their sex: hermaphrodites were identified by their ability to produce offspring in the absence of a mating partner, females by the lack of progeny, and males by their blunt tails (Kanzaki et al., 2017). We calculated sex percentages based only on non-male progeny (hermaphrodites or females). This is because males are not determined by environmental cues, but by sex chromosome number. Raw data used to calculate sex percentages are at https://figshare.com/s/48b14ef15a76acc5405d.

### Assay with conditioned medium and treatment with pharmacological chemical compounds

To induce *A. freiburgensis* hermaphrodite offspring, the parent hermaphrodites (P0 generation) were incubated in the presence of conditioned medium (CM) at 20 °C (Zuco et al., 2018). The CM was derived from 2-3 week old *A. freiburgensis* liquid cultures (M9 medium with *E. coli* OP50-1). Each P0 was placed at the L4 stage onto a 6 cm plate containing NGM and CM. To simulate high-density conditions, 50 mg of lyophilized CM were dissolved in 200 μl of an overnight culture of OP50-1 and spotted onto the plate. F1 eggs were collected for 3-4 days. Each egg was transferred into a single well of a 48-well microtiter plate containing NGM and OP50-1, but no conditioned medium.

For the pharmacological manipulation of signaling pathways, we added compounds to the NGM and OP50-1. The concentration of the compounds was calculated for the volume of the NGM and OP50-1 used. P0 hermaphrodites were incubated with the compounds for 48-36 h at 20 °C. Information about the providers and catalog number for the compounds used in this study are listed in https://figshare.com/s/48b14ef15a76acc5405d.

Chemical compounds were used at the following concentrations: 100 mM Metformin, 6 mM Phenformin, 1 μM Rotenone, 5 μM Forskolin, 30 μM Fluvastatin, 0.5 mM AICAR, 0.5 mM 8-Br-cAMP, 5 mM Salicylate, 10 μM ZLN204, 30 μM O-304, 1 μM Dorsomorphin, 100 μM Rapamycin, 100 μM Methotrexate, 100 nM VO-OHpic, 20 μM Indole-3-Carbinol, 10 μM SRT1720, 100 μM Trichostatin A, 4 mM Valproic Acid, 5 mM DL-beta hydroxybutyrate, 5 mM sodium butyrate, 100 μM EX-527, 3 mM 4-tert-butylbenzoic acid, 75 nM SC-66, 300 nM Akti-1/2, 20 μM perifosine, and 10 nM SC-79. For nematodes incubated with diluted CM, we used 5 mg of freeze-dried CM dissolved in 200 μl *E. coli* OP50-1.

### Inmunohistochemistry

Hermaphrodites were dissected on a slide (Superfrost microscope slide, VWR) in PBS 1X buffer. Dissected gonads were covered by a coverslip and placed on a frozen metal block at −20 °C for at least 10 minutes, and fixed for 2 minutes in a 95% methanol solution at −20 °C. This was followed by 30 minutes in a fixative solution [PBS 1X, 80 mM HEPES (pH= 7.0-7.4), 1.6 mM MgSO4, 0.8 mM EDTA (pH=8.0), 4% paraformaldehyde] in a humid chamber at room temperature. Slides were washed twice with PBST (PBS + 0.1% Triton X-100) for 5 minutes and blocked in PBST + 0.5% BSA for 45-60 minutes. The source of primary and secondary antibodies, as well as dilutions used, are listed in https://figshare.com/s/48b14ef15a76acc5405d. All antibodies were diluted in PBST. Incubation with the primary antibodies was performed at 4 °C overnight. Slides were then washed twice in PBST for 10 minutes each and the corresponding secondary antibody was added and incubated for 2 hours at room temperature. Slides were washed in PBST as above to remove the excess of the secondary antibody and then one drop of Fluoroshield Mounting Medium with 4’,6-diamidino-2-phenylindole (DAPI) (Abcam, #ab104139) was added on the immunostained samples.

Images were taken with a 60X objective in 2.40 μm z-stack intervals (12 sections) with a DeltaVision microscope (Olympus). Acquisition and constrained iterative deconvolution of the images from DeltaVision were processed using the softWoRx software (Applied Precision). The intensity of fluorescence for the secondary antibodies was measured using the ImageJ software (NIH Image, Bethesda, MD).

### Western blot

Protein extraction and buffer preparation were performed following the protocol of (Jeong et al., 2018). Six hundred adult hermaphrodites were collected for each sample: control (OP50-1 only) and experimental (50 mg conditioned medium powder per 200 μl of OP50-1) samples. Protein concentration was measured using Bradford assay (Bradford Reagent, Bio-Rad). We loaded approximately 100 μg of protein. The primary antibodies, against Phospho-AMPKα (Thr172) and PAR-4/LKB1, were used at 1:1000 dilution. The source of primary and secondary antibodies, as well as dilutions used for them, are listed in https://figshare.com/s/48b14ef15a76acc5405d. To detect the signal for the antibodies, we used the Amersham™ ECL™ Western Blotting Detection Reagents (RPN2209).

### Statistical analyses

Results were presented using the most recent developments in data analysis and presentation (Ho et al., 2019), showing the raw data as ‘bee swarm’ plots. They summarize the data showing the mean and the 95% confidence interval (CI), as well as the sampling error distribution diagrammed as a filled curve. These plots provide transparency of the comparison being made, visual clarity and statistical evaluation of the data.

## Supplemental Figures

**Supplemental Figure 1.**
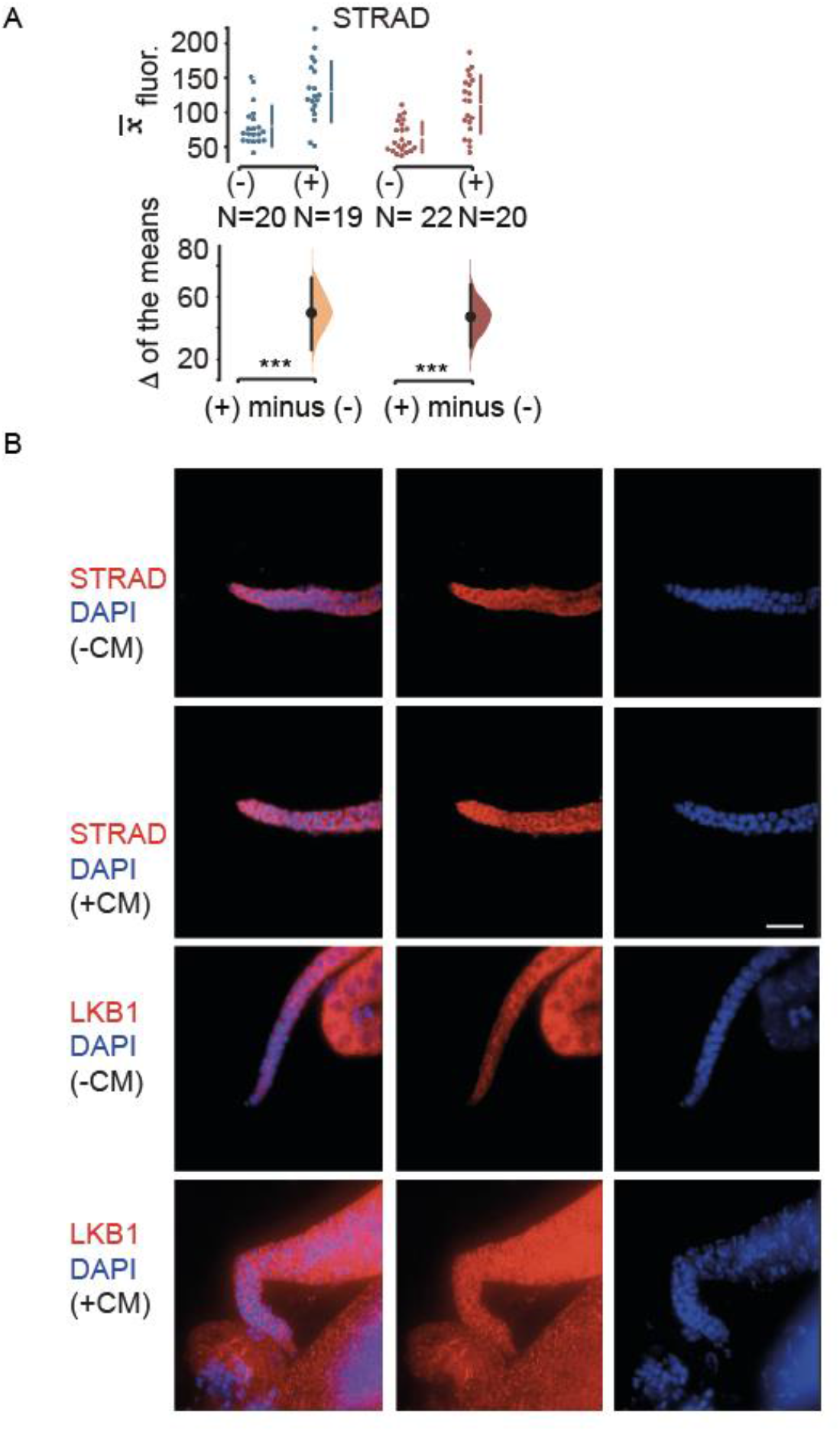
STRAD and LKB1 antibody staining is higher in animals in crowding conditions. **A.** Mean antibody fluorescence (*x__*) in the pre-meiotic (blue) and meiotic portion (red) of the germline, in the absence (-) or presence (+) of conditioned medium. N= sample sizes. Graphical representation as Fig. 2, with ***= p ≤ 0.001. **B.** LKB1 and STRAD in the germline. Staining for antibodies (in red) against LKB1 and STRAD of gonads dissected from hermaphrodites incubated in the presence of either (-) CM or (+) CM. The DNA was stained with DAPI (blue). Bar, 15 μm.

**Supplemental Figure 2.**
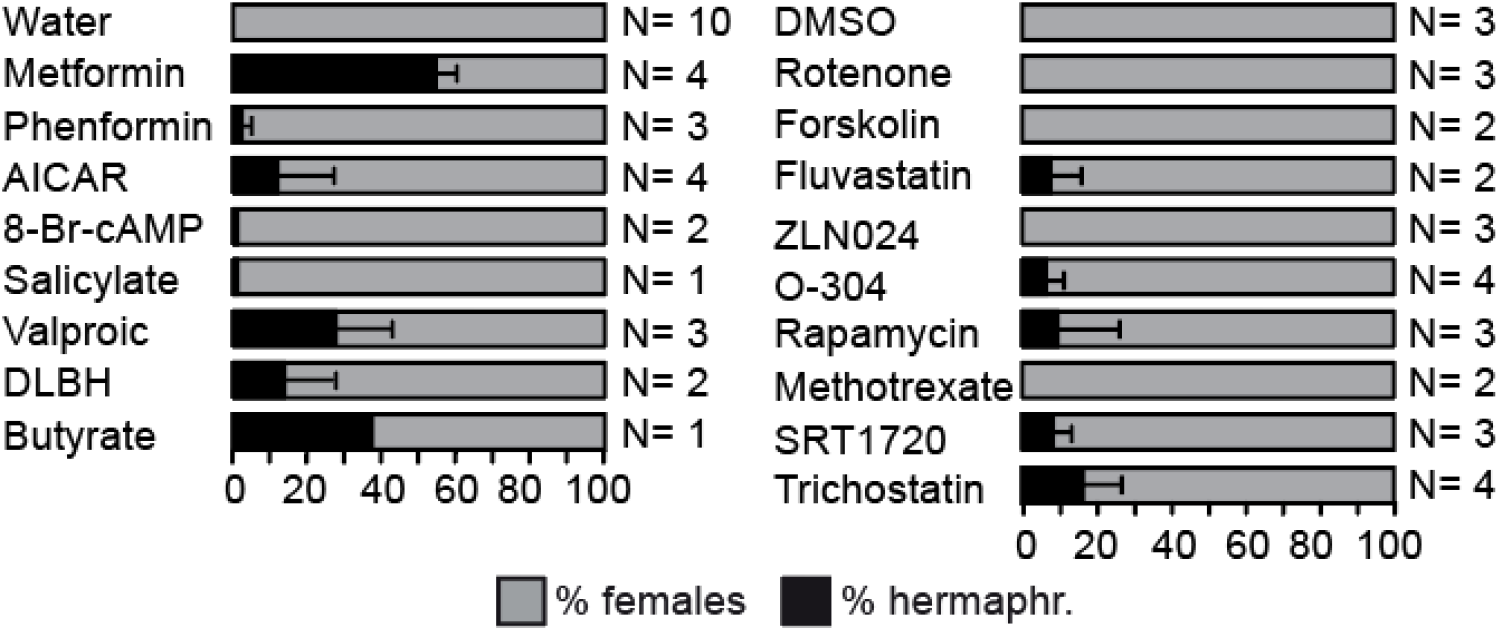
Most chemical compounds affect the sex ratios of hermaphrodites in the absence of diluted CM. Mean percentage and SD of hermaphrodite and female F1 offspring from hermaphrodites treated with chemicals. Chemicals were dissolved either in water or in DMSO. N= number of replicates.

**Supplemental Figure 3.**
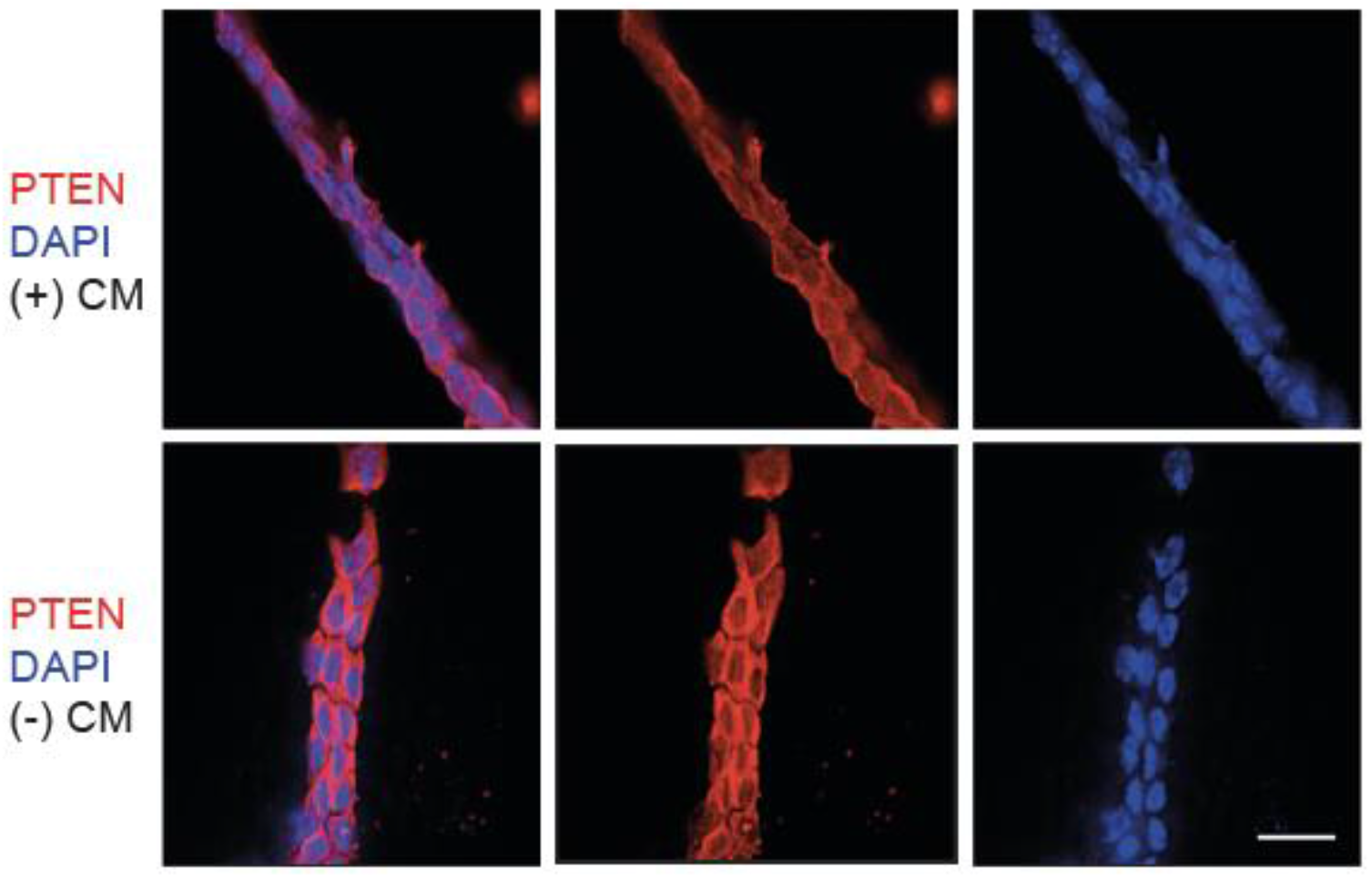
PTEN/DAF-18 in the germline is cytoplasmic and higher in non-crowding conditions. Staining for antibodies (in red) against PTEN/DAF-18 of gonads dissected from hermaphrodites incubated in the presence of either (-) CM or (+) CM. The DNA was stained with DAPI (blue). Bar, 15 μm.

**Supplemental Figure 4.**
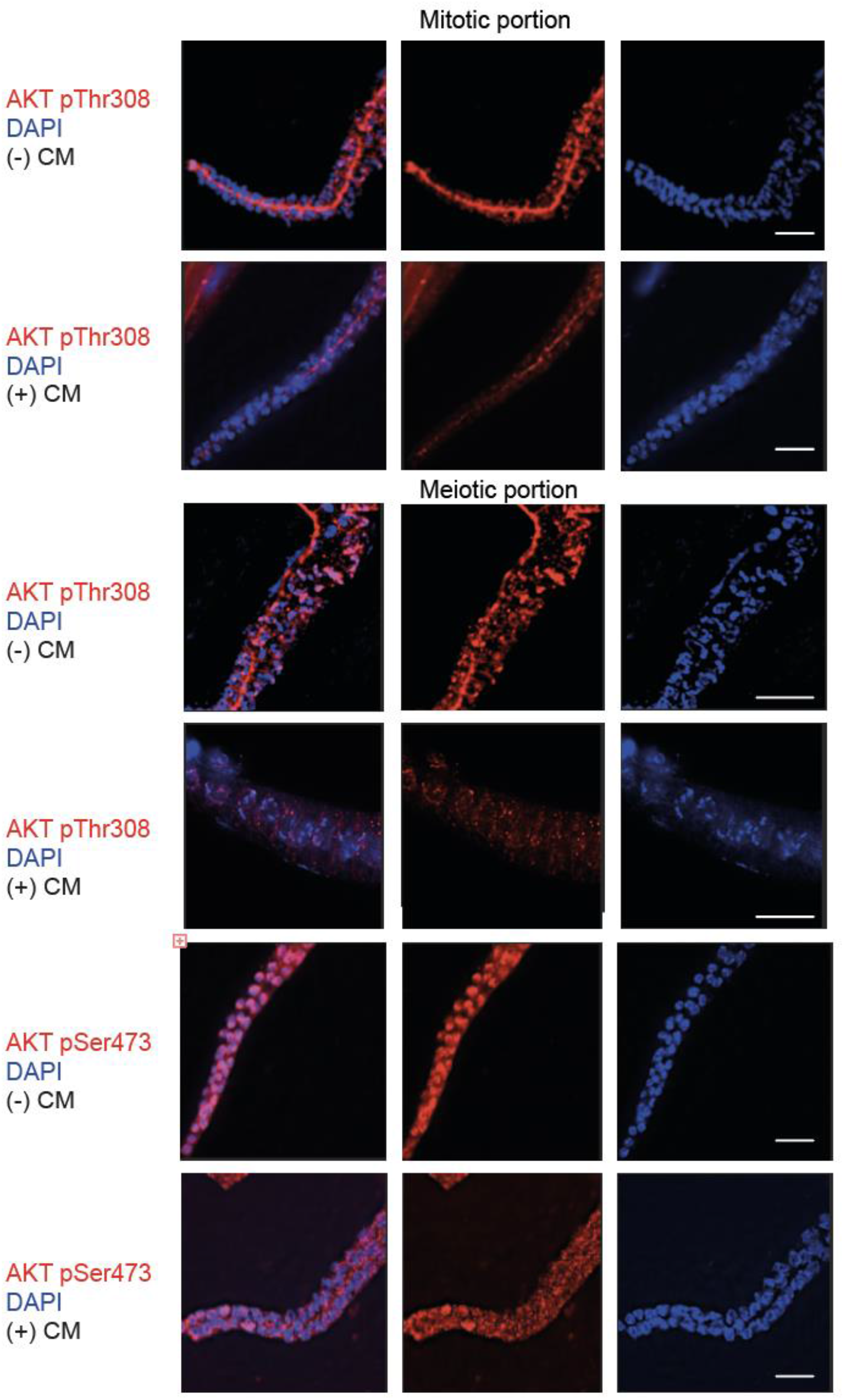
AKT pThr308 in the germline is nuclear and higher in crowding conditions. Staining for antibodies (in red) against AKT pThr308 of gonads dissected from hermaphrodites incubated in the presence of either (-) CM or (+) CM. The DNA was stained with DAPI (blue). Bar, 15 μm.

**Supplemental Figure 5.**
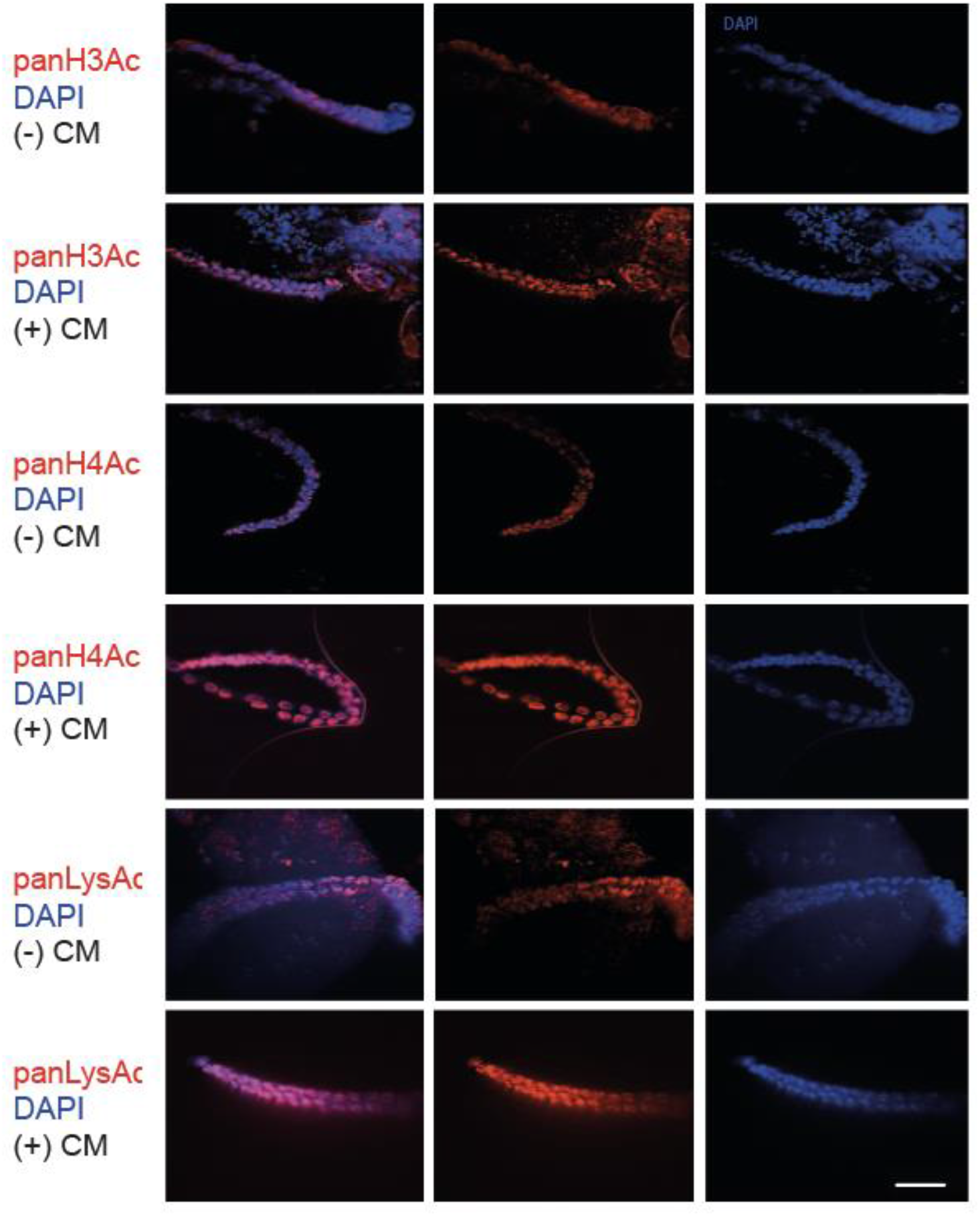
Acetylation in the germline is nuclear and higher in crowding conditions. Staining for antibodies (in red) against panH3Ac, panH4Ac and panLysAc of gonads dissected from hermaphrodites incubated in the presence of either (-) CM or (+) CM. The DNA was stained with DAPI (blue). Bar, 15 μm.

